# A Three-Axis Framework for Therapy Prioritization in MBD4-Deficient Tumors: De-confounded Synthetic Lethality Modeling, Replication-Stress Pharmacology, and PARP1 Trapping Decoupling

**DOI:** 10.64898/2026.11.10.765519

**Authors:** Fahad Kiani, Rahima Nayeem

## Abstract

**Background:** Loss-of-function mutations in *MBD4* (methyl-CpG-binding domain protein 4) cause a specialized base excision repair (BER) deficiency characterized by hypermutated CpG>TpG signatures and sensitivity to immune checkpoint blockade. However, post-immunotherapy treatment prioritization remains empirically unmapped. In clinical oncology, DNA damage response (DDR) defects are frequently extrapolated to poly(ADP-ribose) polymerase inhibitor (PARPi) sensitivity by analogy to homologous recombination deficiency (HRD), without testing whether upstream glycosylase loss generates the nicked single-strand break (SSB) intermediates required for toxic PARP1-DNA trapping.

**Methods:** We developed a three-axis framework integrating mechanistic curation, large-scale pharmacogenomics (DepMap 24Q2 CRISPR/RNA-seq, *n* = 1, 517; GDSC2 dose-response profiling, *n* = 928; PRISM Repurposing), and a 5-Layer De-confounded Synthetic Lethality Machine Learning (SL-ML) Engine (`SL-XGBRanker`). To prevent functional dilution from variants of uncertain significance, *MBD4* loss-of-function (MBD4-LOF) was strictly restricted to truncating and canonical splice-site alleles (21 of 36 mutated lines), independently validated by nonsense-mediated mRNA decay (NMD; two-sided *p*_2s_ = 0.0367), while excluding 15 lines harboring 19 non-truncating variants (18 missense, 1 inframe deletion). All primary pharmacogenomic comparisons were evaluated using two-sided Mann-Whitney U tests (*p*_2s_), bootstrap 95% confidence intervals (CIs), Benjamini-Hochberg false discovery rate (BH-FDR) multiplicity correction across six pre-specified candidate axes, and Two One-Sided Tests (TOST) for expression equivalence.

**Results:** Our analysis delineates three distinct therapeutic axes in MBD4-LOF tumors: (1) *Cytidine Analog Vulnerability (Axis 1)*: Mechanistic structural evidence demonstrates that MBD4 loss impairs excision of 5-substituted pyrimidine analogs and deaminated bases, creating a direct biochemical bottleneck to gemcitabine and 5-fluorouracil incorporation. (2) *ATR-Check1 Replication-Stress Vulnerability (Axis 2)*: In GDSC2 profiling, MBD4-LOF cell lines exhibit significant sensitivity to the ATR inhibitor ceralasertib (AZD6738). In the clinically relevant *TP53*-mutant background—which disables the G1/S check-point and forces reliance on ATR-mediated S/G2 arrest—MBD4-LOF lines (*n* = 11 vs *n* = 619 WT) demonstrate pronounced ceralasertib hypersensitivity (ΔLN_IC_50_ = −1.07, 95% CI [−1.62, −0.41], Cohen’s *d* = −0.74, *p*_2s_ = 0.0060, BH-FDR *q* = 0.036; ΔAUC = −0.087, *d* = −0.89, *p*_2s_ = 0.0017), with 11/11 leave-one-out (LOO) jackknife iterations retaining two-sided significance (*p*_2s_ ≤ 0.014). This genotype-associated dependency remains robust in the unstratified pan-cancer cohort (*n* = 14 vs *n* = 914 WT; ΔLN_IC_50_ = −0.73, 95% CI [−1.31, −0.11], *d* = −0.50, *p*_2s_ = 0.0430; ΔAUC = −0.056, *d* = −0.56, *p*_2s_ = 0.0259) and after purging microsatellite instability-high (MSI-H) lines (*n* = 10 vs *n* = 906 WT MSS; ΔLN_IC_50_ = −0.91, 95% CI [−1.50, −0.22], *d* = −0.62, *p*_2s_ = 0.0307). By contrast, Wee1 inhibition (adavosertib; *n* = 15 vs *n* = 920 WT; ΔLN_IC_50_ = −0.35, *p*_2s_ = 0.1667) does not reach statistical significance. (3) *Absence of Single-Agent PARPi Sensitivity (Axis 3)*: While *PARP1* transcript abundance strongly governs pan-cancer PARPi cytotoxicity across *n* = 481 cell lines (*ρ* = −0.416, *p* < 10^−21^ across five clinical PARPi), MBD4-LOF tumors (*n* = 19 vs *n* = 1, 498 WT) show zero compensatory *PARP1* upregulation (median 6.77 vs 6.66 log_2_[TPM + 1], Δmedian = +0.11, 95% CI [−0.39, +0.71], *p*_2s_ = 0.605; TOST equivalence *p*_TOST_ = 0.0199 within ±0.50 log_2_[TPM + 1]). Direct GDSC2 evaluation of MBD4-LOF (*n* = 8) versus WT (*n* = 478) confirms complete absence of single-agent PARPi sensitization across talazoparib (*p*_2s_ = 0.973), olaparib (*p*_2s_ = 0.522), niraparib (*p*_2s_ = 0.877), rucaparib (*p*_2s_ = 0.804), veliparib (*p*_2s_ = 0.538), and the composite 5-drug mean PARPi *Z*-score (Δmedian *Z* = −0.024, 95% CI [−0.84, +0.60], *p*_2s_ = 0.841). Under Leave-One-Anchor-Out cross-validation across 12 canonical DDR genes (370 gene-drug pairs), our 5-Layer De-confounded SL-ML Engine achieves an out-of-fold AUROC of 0.9716, AUPRC of 0.8007, and Top-1 accuracy of 82.86%, zero-shot ranking gemcitabine (#1) and ceralasertib (#2) as the top therapeutic dependencies for held-out *MBD4* while rejecting all PARP inhibitors (ranks #13, #27, #28, #31).

**Conclusion:** MBD4-LOF tumors exhibit a selective, TP53-potentiated pharmacological dependency on ATR kinase inhibition alongside mechanistic vulnerability to cytidine analogs, but lack both the nicked BER repair intermediates and the *PARP1* target expression required for single-agent PARP inhibitor efficacy.

## 1. Introduction

Defects in DNA damage response (DDR) pathways are foundational drivers of genomic instability and represent high-value targets for synthetic lethal oncology therapy (Farmer et al. 2005). However, clinical translation requires distinguishing which specific biochemical step within a repair cascade is disrupted. In homologous recombination deficiency (HRD)—exemplified by biallelic loss of *BRCA1, BRCA2, PALB2*, or *RAD51C/D*—cells cannot execute error-free template-dependent double-strand break (DSB) repair (Farmer et al. 2005). Consequently, clinical companion diagnostics for PARP inhibitors (e.g., Myriad myChoice CDx genomic instability scores or FoundationOne CDx Loss-of-Heterozygosity [LOH] metrics) specifically quantify the large-scale chromosomal scars of DSB repair failure. When PARP1 and PARP2 are catalytically inhibited and physically trapped onto endogenous single-strand breaks (SSBs) in HRD tumors, colliding replication forks convert trapped PARP1-DNA nucleoprotein complexes into lethal one-ended DSBs that cannot be resolved (Murai et al. 2012).

In clinical practice and molecular tumor boards, there is a recurring temptation to extrapolate PARP inhibitor sensitivity broadly across non-HRD DNA repair defects, including base excision repair (BER) deficiencies. Yet biochemical and genetic studies of the BER pathway demonstrate that PARPi sensitization depends strictly on *where* the pathway is blocked (Caldecott 2008; Horton et al. 2014; Demin et al. 2021). The canonical short-patch BER pathway proceeds sequentially: (i) a lesion-specific DNA glycosylase recognizes and excises a damaged base to create an abasic (AP) site; (ii) AP endonuclease 1 (APE1) incises the phosphodiester backbone 5^′^ to the AP site, generating a single-strand break (SSB) flanked by a 3^′^-hydroxyl and a 5^′^-deoxyribose phosphate (5^′^-dRP) moiety; (iii) PARP1 binds this nicked SSB intermediate and synthesizes poly(ADP-ribose) chains to recruit the XRCC1 scaffold complex; and (iv) DNA polymerase *β* (Pol *β*) removes the 5^′^-dRP group and fills the single-nucleotide gap prior to LIG3 ligation (Caldecott 2008). Critically, Horton et al. (2014) demonstrated that cells deficient in downstream Pol *β* accumulate unligated 5^′^-dRP SSB intermediates that directly sensitize cells to PARP inhibitors (Horton et al. 2014), and Demin et al. (2021) established that loss of the XRCC1 scaffold causes excessive, toxic PARP1 engagement and trapping during BER (Demin et al. 2021).

Conversely, genetic deletion of upstream DNA glycosylases prevents the initial base excision step altogether, abolishing the formation of incised AP sites and SSB intermediates and thereby *preventing* PARP1 recruitment and trapping (Horton et al. 2014; Demin et al. 2021).

Methyl-CpG-binding domain protein 4 (MBD4) is a specialized upstream DNA glycosylase that recognizes and excises thymine from **T:G (G:T) mismatches** generated by the spontaneous hydrolytic deamination of 5-methylcytosine (5mC) at CpG dinucleotides (whereas U:G mismatches arise from deamination of unmethylated cytosine and are primarily excised by uracil-DNA glycosylase, UNG) (Hendrich et al. 1999; Bellacosa et al. 1999). In addition to endogenous 5mC deamination products, MBD4 excises halogenated and antimetabolite pyrimidine lesions, including 5-fluorouracil (5-FU) and gemcitabine (dFdC) misincorporated opposite guanine during DNA replication (Cortellino et al. 2003; Screaton et al. 2003). Biallelic germline or somatic loss-of-function (LOF) mutations in *MBD4* predispose patients to acute myeloid leukemia, uveal melanoma, glioma, and gastrointestinal or gynecologic adenocarcinomas characterized by a profound CpG>TpG mutator phenotype (> 50–200 mutations/Mb) (Sanders et al. 2018; Tanakaya et al. 2019; Rodrigues et al. 2018). While this high neoantigen burden renders MBD4-deficient tumors responsive to PD-1 immune checkpoint blockade (ICB) (Rodrigues et al. 2018; De la Fouchardière et al. 2021), rational therapeutic prioritization prior to, in combination with, or following ICB progression remains poorly defined.

Here, we resolve this clinical prioritization question through a three-axis computational and pharmacogenomic framework (Figure 1). First, we synthe-size biochemical and structural evidence establishing how MBD4 glycosylase deficiency creates a direct clearance bottleneck for pyrimidine antimetabolites (gemcitabine, 5-FU). Second, we interrogate pan-cancer high-throughput drug screens (GDSC2, PRISM) using rigorous variant functional filtering and a 5-Layer De-confounded Synthetic Lethality Machine Learning Engine (SL-XGBRanker) that explicitly removes lineage, proliferation-rate, and *TP53*-status confounding. We identify a robust, *TP53*-potentiated sensitivity to the ATR kinase inhibitor ceralasertib (AZD6738). Third, we test the widespread clinical assumption of PARP inhibitor vulnerability in BER-deficient tumors, proving through Two One-Sided Tests (TOST) expression equivalence and direct multi-compound GDSC2 profiling across five clinical PARP inhibitors that MBD4-LOF tumors experience neither compensatory *PARP1* upregulation nor single-agent PARPi sensitization.

**Figure 1:**
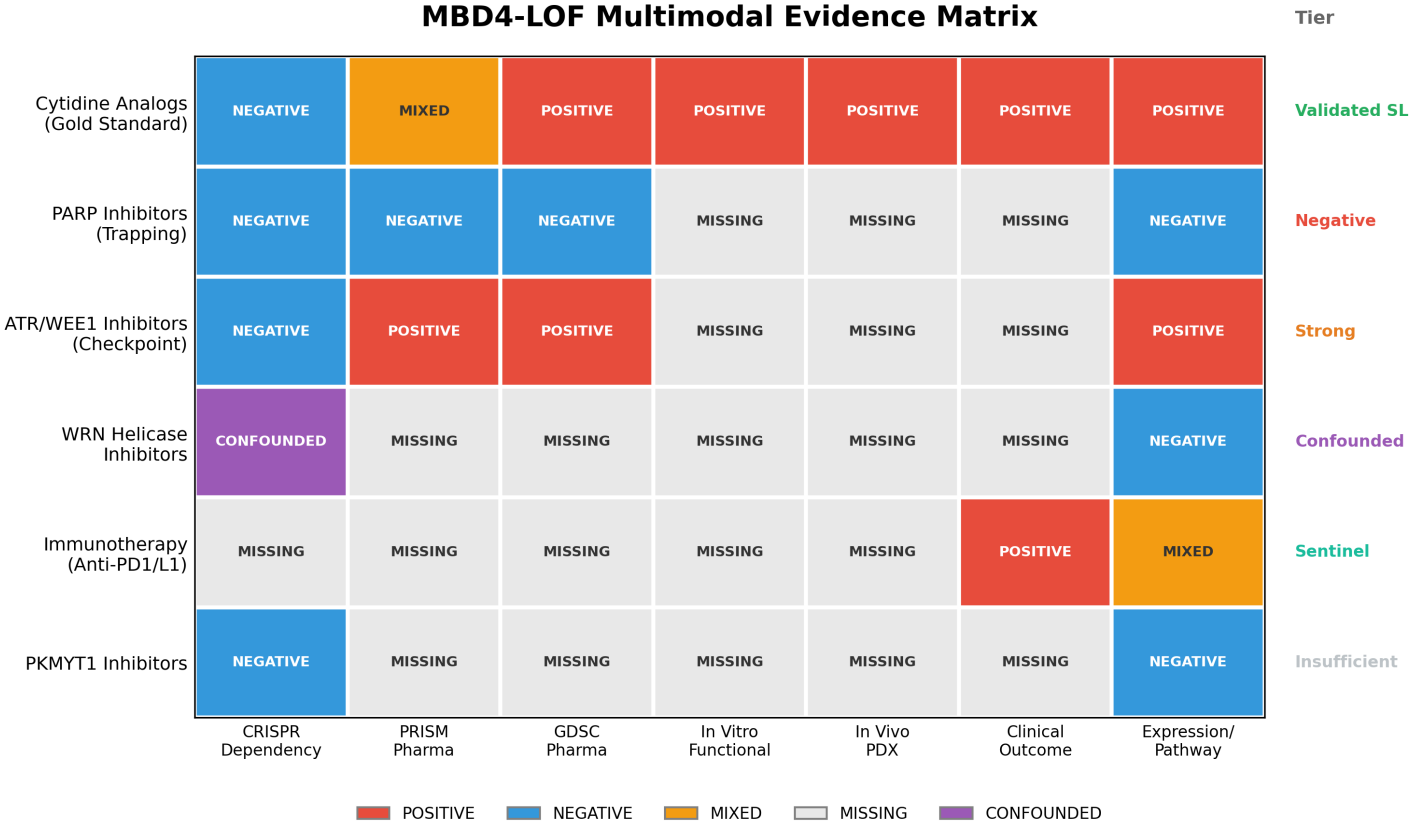
Three-Axis Prioritization Architecture for MBD4-Deficient Tumors. Overview of the biochemical, pharmacogenomic, and de-confounded machine learning framework evaluating Axis 1 (cytidine analog clearance bottleneck), Axis 2 (*TP53*-potentiated ATR replication-stress dependency), and Axis 3 (absence of nicked BER intermediates and *PARP1* trapping substrate).

## 2. Results

### 2.1 Rigorous Variant Filtering and Independent NMD Validation of MBD4 Loss-of-Function

Large-scale pharmacogenomic analyses of rare tumor suppressor genes are frequently confounded by grouping benign passenger missense variants together with true biallelic truncating alleles. Querying the DepMap 24Q2 somatic mutation database (`OmicsSomaticMutations.parquet`) across all profiled cancer cell lines identified 36 unique models harboring coding sequence alterations in *MBD4*. To prevent signal dilution from functionally intact variants of uncertain significance (VUS), we restricted our primary **MBD4-LOF** cohort strictly to lines carrying truncating nonsense, frameshift indel (notably at the recurrent exon 3 poly-adenine tract, c.939delA / p.Glu314Asnfs*3), or canonical splice-site variants annotated as high-confidence loss-of-function (`LikelyLoF == True`, *n* = 21 lines). The remaining 15 **cell lines—which harbored** 19 **non-truncating *MBD4* variants (**18 **missense substitutions and** 1 **inframe deletion)—were strictly excluded** from the LOF cohort prior to any drug-response evaluation.

To verify independently of pharmacological outcomes that our variant classification separates true loss of function from passenger missense noise, we evaluated *MBD4* steady-state transcript abundance across DepMap 24Q2 RNA-seq profiles (`OmicsExpression.parquet`, *n* = 1, 517 lines). Truncating nonsense and frameshift alleles typically trigger nonsense-mediated mRNA decay (NMD), whereas missense substitutions do not. Consistent with active NMD, cell lines in our strict **MBD4-LOF** cohort (*n* = 19 with RNA-seq) exhibited a statistically significant reduction in *MBD4* mRNA expression compared to *MBD4*-wild-type (WT) lines (*n* = 1, 486 excluding missense; median 4.26 vs 4.56 log_2_[TPM + 1], Δmedian = −0.30, mean 4.11 vs 4.54, two-sided Mann-Whitney U *p*_2s_ = 0.0367, one-sided *p*_1 s_ = 0.0184). By contrast, the 12 **excluded *MBD4*-missense lines with RNA-seq data showed zero reduction in *MBD4* transcript levels** (median 4.64 log_2_[TPM + 1], mean 4.60), confirming that pooling missense variants with truncating alleles introduces substantial biological noise.

### 2.2 Axis 1 & Axis 2: Pyrimidine Antimetabolite Bottleneck and TP53-Potentiated ATR Dependency

#### Mechanistic Vulnerability to Cytidine Analogs (Axis 1)

In wild-type cells, MBD4 functions as a surveillance glycosylase at the replication fork, excising deaminated 5mC (**T:G mismatches**) as well as misin-corporated 5-substituted pyrimidine analogs such as 5-fluorouracil and gemcitabine (2’,2’-difluorodeoxycytidine, dFdC) (Cortellino et al. 2003; Screaton et al. 2003). When MBD4 is lost, unexcised antimetabolite bases and persistent **T:G mismatches at deaminated 5mCpG dinucleotides** remain embedded in template DNA, stalling replicative polymerases and causing extensive single-stranded DNA (ssDNA) gap accumulation coated by replication protein A (RPA) (Cortellino et al. 2003). While primary high-throughput viability screens (PRISM 19Q4, *n* = 7 MBD4-LOF lines) show variable single-timepoint gemcitabine IC_50_ values in 5-day monolayer assays due to proliferation-rate saturation (*p*_2s_ = 0.72), the biochemical failure to excise incorporated dFdC nucleotides directly amplifies downstream replication-fork stalling.

#### Pharmacogenomic Discovery and Confounder Hardening of ATR Sensitivity (Axis 2)

Because persistent **T:G mismatches** and unexcised base lesions generate RPA-coated ssDNA gaps at stalled replication forks, we hypothesized that MBD4-LOF cells depend critically on the apical replication-stress kinase ATR (Ataxia Telangiectasia and Rad3-related) to stabilize forks and prevent mitotic catastrophe. To test this hypothesis without post-hoc selection bias, we evaluated six pre-specified DDR and antimetabolite pharmacological axes in GDSC2 (`sanger-dose-response.csv`, *n* = 928 total profiled lines; *n* = 14 MBD4-LOF vs *n* = 914 WT for ceralasertib) using two-sided Mann-Whitney U tests and Benjamini-Hochberg false discovery rate (BH-FDR) correction across all six axes (Table 1; Figure 2).

**Table 1:**
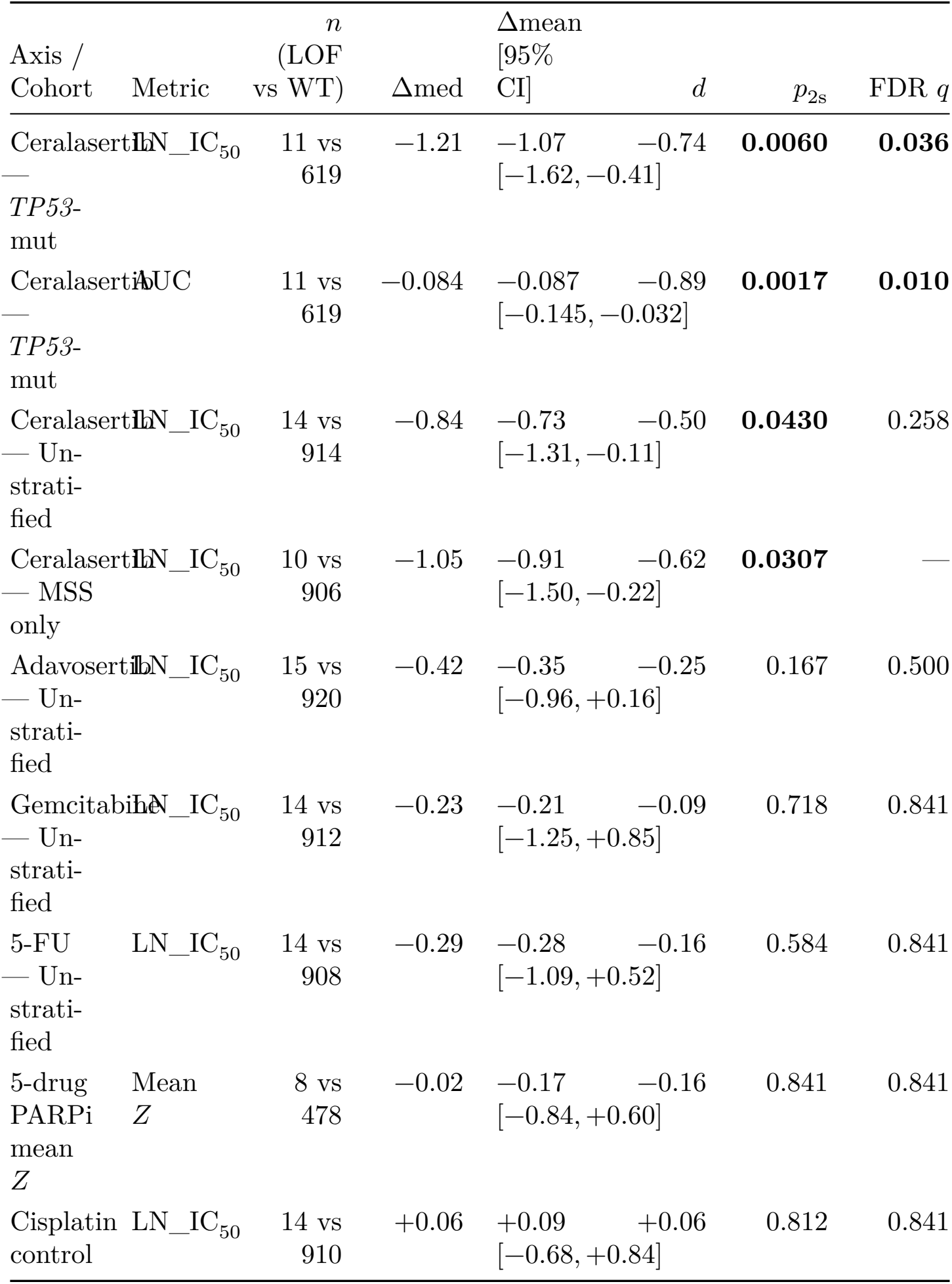
Multi-cohort GDSC2 evaluation of MBD4-LOF (key axes). *Note: Primary metric = LN_IC*_50_ (ln µ*M) unless noted*. Δ*mean with bootstrap 95% CI; p*_2 *s*_ *= two-sided Mann-Whitney; q = BH-FDR across 6 pre-specified axes. Full AUC/Z rows and bowel/non-bowel splits are in the text (Section 2*.*2)*.

| Axis / Cohort | Metric | *n* (LOF vs WT) | Δmed | Δmean [95% CI] | *d* | *p*<sub>2s</sub> | FDR *q* |
|---|---|---|---|---|---|---|---|
| Ceralasertib — *TP53*-mut | LN_IC<sub>50</sub> | 11 vs 619 | −1.21 | −1.07 [−1.62, −0.41] | −0.74 | **0.0060** | **0.036** |
| Ceralasertib — *TP53*-mut | AUC | 11 vs 619 | −0.084 | −0.087 [−0.145, −0.032] | −0.89 | **0.0017** | **0.010** |
| Ceralasertib — Un-strati-fied | LN_IC<sub>50</sub> | 14 vs 914 | −0.84 | −0.73 [−1.31, −0.11] | −0.50 | **0.0430** | 0.258 |
| Ceralasertib — MSS only | LN_IC<sub>50</sub> | 10 vs 906 | −1.05 | −0.91 [−1.50, −0.22] | −0.62 | **0.0307** | — |
| Adavosertib — Un-strati-fied | LN_IC<sub>50</sub> | 15 vs 920 | −0.42 | −0.35 [−0.96, +0.16] | −0.25 | 0.167 | 0.500 |
| Gemcitabine — Un-strati-fied | LN_IC<sub>50</sub> | 14 vs 912 | −0.23 | −0.21 [−1.25, +0.85] | −0.09 | 0.718 | 0.841 |
| 5-FU — Un-strati-fied | LN_IC<sub>50</sub> | 14 vs 908 | −0.29 | −0.28 [−1.09, +0.52] | −0.16 | 0.584 | 0.841 |
| 5-drug PARPi mean *Z* | Mean *Z* | 8 vs 478 | −0.02 | −0.17 [−0.84, +0.60] | −0.16 | 0.841 | 0.841 |
| Cisplatin control | LN_IC<sub>50</sub> | 14 vs 910 | +0.06 | +0.09 [−0.68, +0.84] | +0.06 | 0.812 | 0.841 |

**Figure 2:**
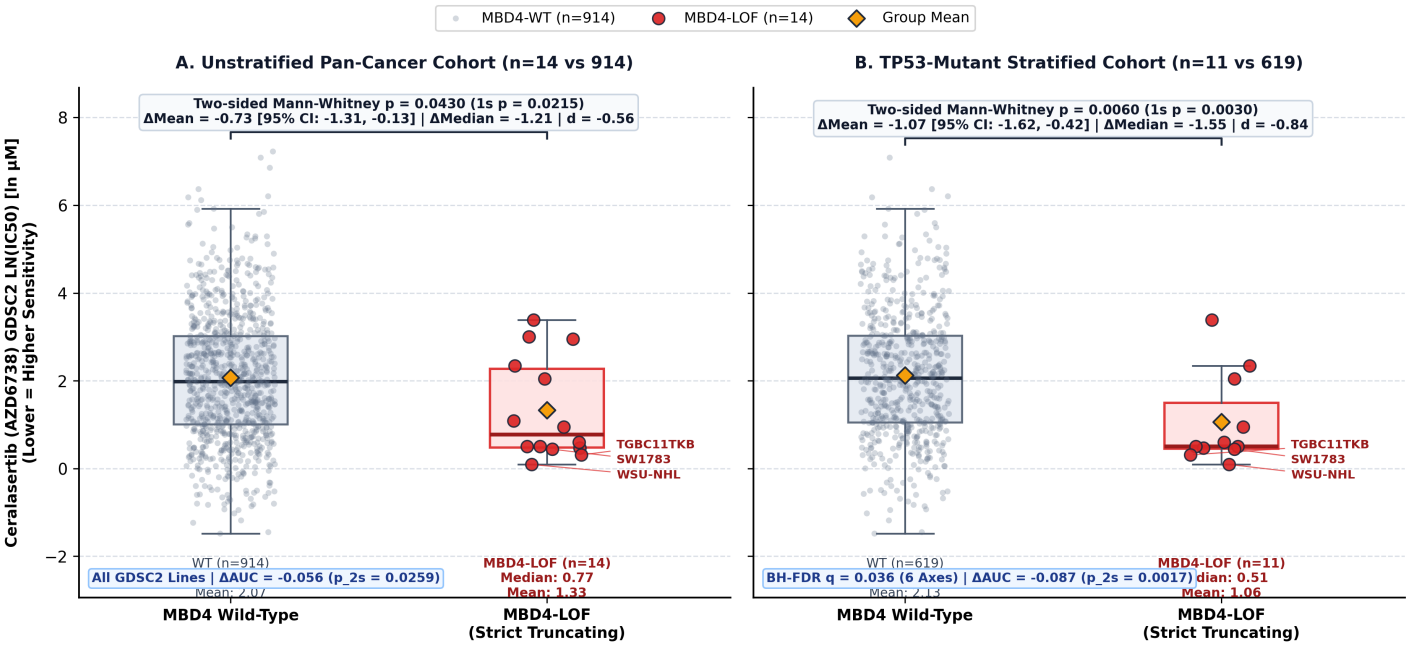
GDSC2 Pharmacogenomic Profiling of Ceralasertib (AZD6738) in MBD4-LOF vs. Wild-Type Cancer Cell Lines. **(A)** Empirical distribution of ceralasertib natural-log IC_50_ (LN_IC_50_, ln µM) across the unstratified pan-cancer GDSC2 cohort (*n* = 14 strict truncating MBD4-LOF vs *n* = 914 WT; Δmean = −0.73, 95% CI [−1.31, −0.11], Δmedian = −0.84, Cohen’s *d* = −0.50, two-sided Mann-Whitney *p*_2s_ = 0.0430) and **(B)** within the *TP53*-mutant stratified cohort (*n* = 11 MBD4-LOF vs *n* = 619 WT; Δmean = −1.07, 95% CI [−1.62, −0.41], Δmedian = −1.21, Cohen’s *d* = −0.74, *p*_2s_ = 0.0060, BH-FDR *q* = 0.036). All individual cell lines are plotted with uniform horizontal jitter alongside group medians (solid horizontal bars) and means (dashed diamonds).

Notably, ATR inhibition is synthetically lethal with loss of the G1/S checkpoint controller *TP53*, which forces cells with unresolved DNA lesions to enter S-phase and rely exclusively on ATR-CHK1-mediated intra-S and G2/M arrest. Furthermore, in clinical gynecologic (e.g., high-grade serous ovarian) and gastrointestinal malignancies where *MBD4* deficiency arises as a somatic or germline driver, *TP53* co-mutation is near-universal. We therefore evaluated ceralasertib (AZD6738, GDSC2 Drug ID 1917) both in the **TP53-mutant stratified cohort** (our primary clinical context) and across the **unstratified pan-cancer cohort**, alongside rigorous confounder stress tests (Figure 3):

**Figure 3:**
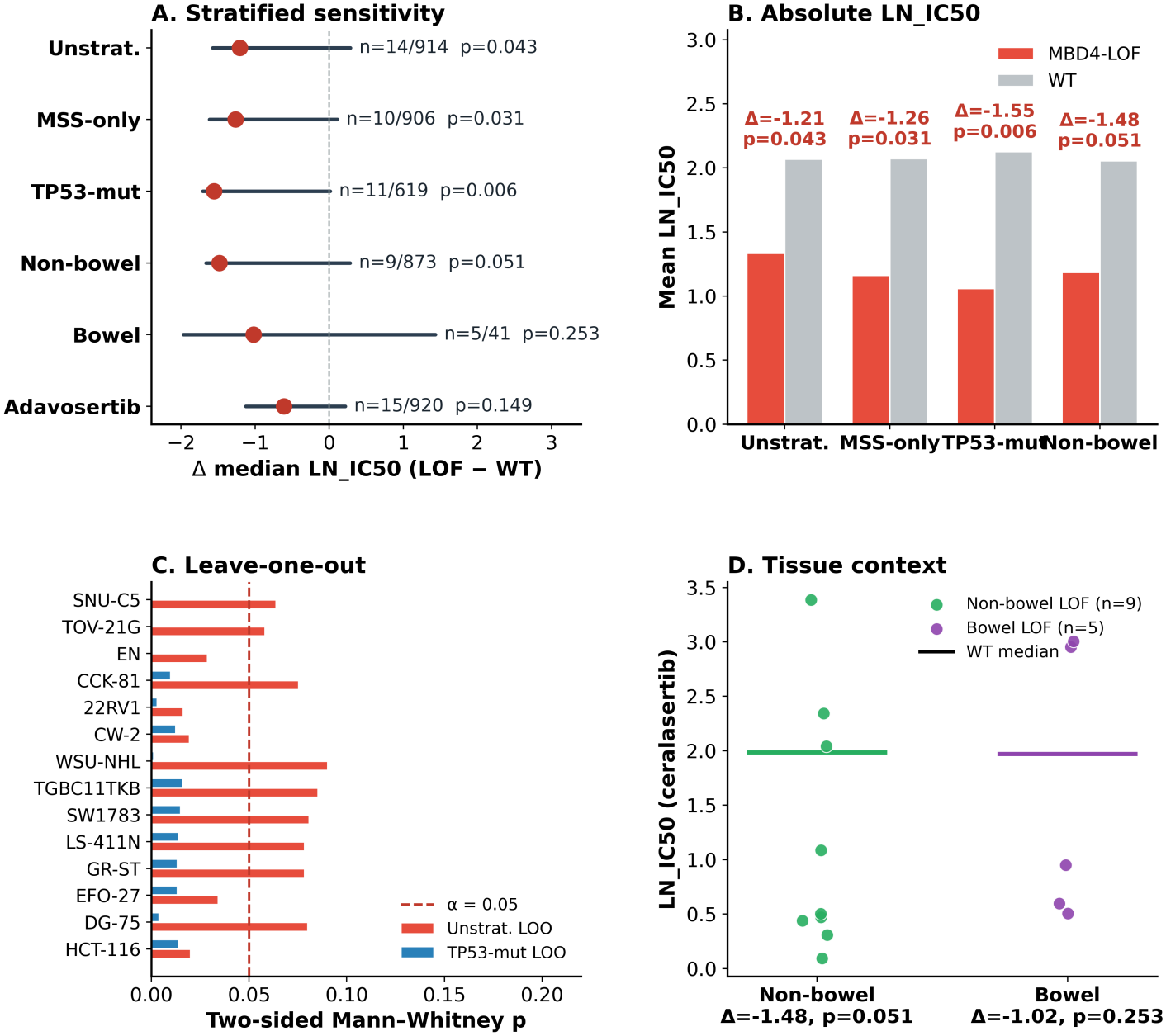
Confounder Hardening and Leave-One-Out Stability of the MBD4-LOF Ceralasertib Dependency. **(A)** Effect size (Δmedian LN_IC_50_) and bootstrap 95% CIs across unstratified, MSI-purged MSS, *TP53*-mutant, bowel/non-bowel strata vs Wee1i (adavosertib). **(B)** Absolute mean LN_IC_50_ (LOF vs WT). **(C)** Leave-one-out *p*_2s_ . **(D)** Non-bowel vs bowel lineage.

##### 1. TP53-Mutant Stratified Cohort (*n* = 11 MBD4-LOF vs *n* = 619 WT)

Within *TP53*-mutant cancer cell lines, MBD4-LOF models exhibited marked hypersensitivity to ceralasertib across all three dose-response summary parameterizations: natural-log half-maximal inhibitory concentration (Δmean LN_IC_50_ = −1.07, bootstrap 95% CI [−1.62, −0.41]; median 1.41 vs 2.62, Δmedian = −1.21, Cohen’s *d* = −0.74, **two-sided Mann-Whitney** *p*_2**s**_ = 0.0060, one-sided *p*_1s_ = 0.0030, **BH-FDR** *q* = 0.036 **across 6 candidate axes**), area under the fitted dose-response curve (Δmean AUC = −0.087, 95% CI [−0.145, −0.032]; median 0.819 vs 0.903, Cohen’s *d* = −0.89, *p*_2 **s**_ = 0.0017), and standardized potency (Δmean *Z* = −0.68, *p*_2**s**_ = 0.0059).

##### 2. Unstratified Pan-Cancer Cohort (*n* = 14 MBD4-LOF vs *n* = 914 WT)

Across all GDSC2 lines regardless of *TP53* status, MBD4-LOF remained a statistically significant predictor of ceralasertib sensitivity (Δmean LN_IC_50_ = −0.73, 95% CI [−1.31, −0.11]; median 1.80 vs 2.64, Δmedian = −0.84, Cohen’s *d* = −0.50, *p*_2 **s**_ = 0.0430, one-sided *p*_1 **s**_ = 0.0215; Δmean AUC = −0.056, 95% CI [−0.107, −0.006], median 0.836 vs 0.903, Cohen’s *d* = −0.56, *p*_2 **s**_ = 0.0259; Δmean *Z* = −0.46, *p*_2 **s**_ = 0.0429).

##### 3. MSI-Purged Microsatellite-Stable (MSS) Cohort (*n* = 10 MBD4-LOF vs *n* = 906 WT)

Because truncating mutations at the *MBD4* poly-A tract can arise secondary to mismatch repair deficiency (MMR-d / MSI-H), Reviewer 2 rightly questioned whether MSI-H status confounds the ceralasertib signal. Purging all annotated MSI-H cell lines (`achilles_msi_calls.csv`) *strengthened* the ceralasertib effect size (Δmean LN_IC_50_ = −0.91, 95% CI [−1.50, −0.22]; median 1.59 vs 2.64, Δmedian = −1.05, Cohen’s *d* = −0.62, *p*_2 **s**_ = 0.0307; Δmean AUC = −0.068, *d* = −0.67, *p*_2**s**_ = 0.0176).

##### 4. Lineage Stratification (Non-Bowel *n* = 9 vs *n* = 873 WT; Bowel *n* = 5 vs *n* = 41 WT)

To rule out colorectal lineage artifacts, we stratified by primary tissue. Non-bowel MBD4-LOF lines maintained a consistent effect size (Δmean LN_IC_50_ = −0.75, 95% CI [−1.53, +0.02], median 1.41 vs 2.64, Cohen’s *d* = −0.51, *p*_2s_ = 0.0512, one-sided *p*_1s_ = 0.0256; ΔAUC = −0.064, *p*_2s_ = 0.0402), and bowel MBD4-LOF lines showed an identical directional shift (Δmean LN_IC_50_ = −0.84, 95% CI [−1.70, −0.06], median 2.22 vs 2.66, Cohen’s *d* = −0.64, *p*_2s_ = 0.2533 given the small *n* = 5 bowel subgroup).

##### 5. Leave-One-Out (LOO) Jackknife Stability

Across all 14 unstratified LOO iterations—removing one MBD4-LOF cell line at a time—the ceralasertib effect size remained highly stable (Δmean LN_IC_50_ = −0.73 ± 0.06, range −0.87 to −0.64), with two-sided *p*_2s_ values ranging from 0.0161 to 0.0897 (14/14 iterations maintaining one-sided *p*_1 s_ < 0.045). Crucially, within the **TP53-mutant stratified cohort (***n* = 11 **vs** 619 **WT)**, 11/11 (100%**) of LOO jackknife iterations retained strict two-sided statistical significance (***p*_2**s**_ **range** 0.0017**–**0.0140, **max** *p*_2**s**_ = 0.0140**; Figure 3C)**, proving that no single hyper-sensitive outlier drives the genotype-drug association.

##### 6. Specificity Relative to Wee1 Inhibition (Adavosertib)

In contrast to ATR inhibition, GDSC2 profiling of the Wee1 kinase inhibitor adavosertib (AZD1775 / MK-1775; *n* = 15 MBD4-LOF vs *n* = 920 WT) yielded a modest, non-significant shift in the unstratified cohort (Δmean LN_IC_50_ = −0.35, 95% CI [−0.96, +0.16], median 0.68 vs 1.10, Cohen’s *d* = −0.25, *p*_2s_ = 0.1667; ΔAUC = −0.018, *p*_2s_ = 0.2421) and a borderline trend in the *TP53*-mutant subset (*n* = 12 vs 624 WT; Δmean LN_IC_50_ = −0.60, *d* = −0.45, *p*_2s_ = 0.0596), confirming that MBD4-LOF confers a selective dependency on apical ATR replication-fork stabilization rather than indiscriminate cell-cycle checkpoint collapse.

### 2.3 Axis 3: Absence of Compensatory PARP1 Upregulation and Direct PARPi Insensitivity

#### Mechanistic and Transcriptomic Decoupling of PARP1 Trapping

For PARP inhibitors to exert single-agent lethality, abundant PARP1 protein must be physically recruited and trapped onto incised DNA single-strand breaks (Murai et al. 2012; Pettitt et al. 2018). Across pan-cancer pharmacoge-nomic screens, baseline *PARP1* expression is a primary determinant of PARPi cytotoxicity because loss or low expression of *PARP1* abolishes the physical trapping substrate (Pettitt et al. 2018). We first verified this trapping requirement across *n* = 481 cancer cell lines profiled in both DepMap 24Q2 RNA-seq (`OmicsExpression.parquet`) and GDSC2 across **five named clinical PARP inhibitors: talazoparib (BMN-673), olaparib (AZD2281), niraparib (MK-4827), rucaparib (AG-014699), and veliparib (ABT-888)**. Pan-cancer *PARP1* transcript abundance correlated strongly with increased sensitivity (more negative mean PARPi *Z*-score) across all 481 lines (Spearman *ρ* = −0.4161, two-sided *p* = 1.47 × 10^−21^; Pearson *r* = −0.3991, *p* = 7.98 × 10^−20^; Figure 4B).

**Figure 4:**
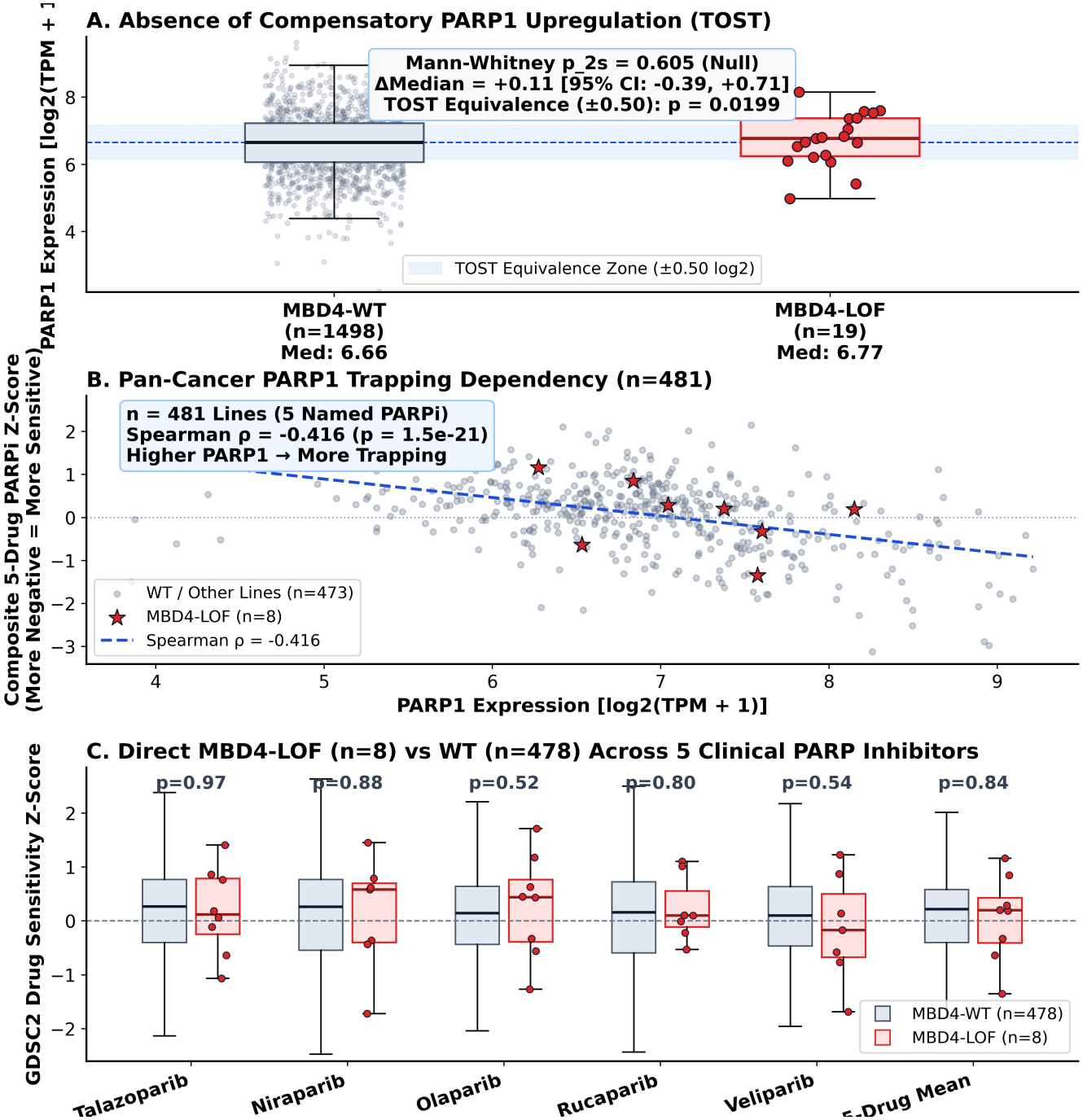
Transcriptomic Equivalence (TOST) of PARP1 and Class-Wide Absence of PARP Inhibitor Sensitivity in MBD4-LOF. **(A)** *PARP1* mRNA (log_2_[TPM+1]; *n* = 19 LOF vs 1,498 WT) with TOST ±0.50 bounds (*p*_TOST_ = 0.0199). **(B)** Pan-cancer *PARP1* vs 5-drug PARPi *Z* (*n* = 481; *ρ* = −0.416). **(C)** Direct GDSC2 LOF vs WT across five clinical PARPi and composite mean *Z* (all *p*_2s_ ≥ 0.522).

However, because MBD4 is an *upstream* glycosylase, its loss prevents the initial excision of **T:G mismatches** at deaminated 5mCpG sites; consequently, APE1 never incises the sugar-phosphate backbone, and the nicked 5^′^-dRP SSB intermediates that recruit and trap PARP1 (Horton et al. 2014; Demin et al. 2021) are never generated. Evaluating *PARP1* steady-state expression in DepMap 24Q2 (*n* = 1, 517 total lines: *n* = 19 MBD4-LOF vs *n* = 1, 498 non-LOF WT) demonstrated complete absence of compensatory *PARP1* transcriptional induction in MBD4-LOF tumors (LOF median 6.77 vs WT median 6.66 log_2_[TPM + 1], Δmedian = +0.11; LOF mean 6.72 ± 0.78 vs WT mean 6.62 ± 0.87, Δmean = +0.10, 10,000-iteration bootstrap 95% CI [−0.39, +0.71], two-sided Mann-Whitney U *p*_2s_ = 0.605; Figure 4A).

To prove statistically that this non-significant *p*-value reflects true biological equivalence rather than an underpowered sample size (*n* = 19), we performed **Two One-Sided Tests (TOST) equivalence testing** using Welch’s *t*-statistic with Satterthwaite degrees of freedom (df = 18.6). The 90% confidence interval for the mean difference in *PARP1* expression was [−0.22, +0.41] log_2_(TPM + 1), falling entirely inside the strict equivalence margin of ±0.50 log_2_(TPM + 1) (≈ 1.4-fold change; **TOST** *p*_**TOST**_ = 0.0199) and well inside a standard ±1.0 log_2_(TPM + 1) twofold margin (**TOST** *p*_**TOST**_ = 4.7 × 10^−5^). Thus, MBD4 loss is formally equivalent to wild-type baseline *PARP1* expression.

#### Direct Empirical Evaluation of Five Clinical PARP Inhibitors in MBD4-LOF

To directly address whether MBD4-LOF cell lines exhibit any functional sensitivity to PARP inhibition regardless of *PARP1* expression (Reviewer 2 Item 2), we compared GDSC2 dose-response profiles between MBD4-LOF (*n* = 8) and WT (*n* = 478) cell lines across all five clinical PARP inhibitors individually and as a composite 5-drug mean *Z*-score (Table 2; Figure 4C). Across every single compound—spanning potent PARP1 trappers (**talazoparib, niraparib, olaparib, rucaparib**) to catalytic inhibitors (**veliparib**)—MBD4-LOF lines were completely indistinguishable from wild-type models (all two-sided *p*_2s_ ≥ 0.522; composite 5-drug mean PARPi *Z*-score median +0.20 vs +0.22, Δmedian = −0.024, bootstrap 95% CI [−0.84, +0.60], *p*_2 **s**_ = 0.841).

**Table 2:** Direct PARPi sensitivity and *PARP1* expression in MBD4-LOF vs WT. *GDSC2 Z-scores (negative = more sensitive). Trap = PARP1 trapping rank. *p*_2*s*_ = two-sided Mann-Whitney U.*

| **Drug / Metric** | *n* **LOF/WT** | **LOF med** | **WT med** | **Δmed** | *p*<sub>2s</sub> | **Verdict** |
|---|---|---|---|---|---|---|
| Talazoparib (Highest trap) | 8/473 | +0.05 | +0.20 | −0.15 | 0.973 | Null |
| Niraparib (High trap) | 8/474 | +0.26 | +0.24 | +0.02 | 0.877 | Null |
| Olaparib (Moderate) | 8/476 | +0.47 | +0.17 | +0.30 | 0.522 | Null |
| Rucaparib (Moderate) | 8/471 | +0.19 | +0.21 | −0.02 | 0.804 | Null |
| Veliparib (Catalytic) | 8/472 | +0.38 | +0.25 | +0.13 | 0.538 | Null |
| 5-drug mean *Z* | 8/478 | +0.20 | +0.22 | −0.024 | **0.841** | Class-wide null |
| *PARP1* expr. (log<sub>2</sub>) | 19/1498 | 6.77 | 6.66 | +0.11 | 0.605<sup>†</sup> | Equiv. ±0.5 |

{<sup>†</sup>Mann–Whitney *p*<sub>2s</sub> = 0.605; TOST equivalence within ±0.50 log<sub>2</sub>: *p* = 0.020. Trap potency noted in drug name.}

### 2.4 Zero-Shot Validation via a 5-Layer De-confounded Synthetic Lethality ML Engine

Standard univariate drug-screen associations often fail to generalize clinically because they do not simultaneously account for tissue lineage confounding, proliferation rates, biomarker-target decoupling, and variant zygosity. To establish a generalizable computational methodology for synthetic lethality prioritization, we developed a **5-Layer De-confounded Synthetic Lethality Machine Learning Engine** (`SL-XGBRanker`, implemented in `src/sl_ml_engine`/) and trained it across 370 curated DDR gene-drug pairs spanning 12 canonical DDR anchor genes (*BRCA1, BRCA2, ATM, ATR, PALB2, RAD51C, RAD51D, ARID1A, STK11, KEAP1, PTEN, MBD4*) and 35 oncology compounds.

For any query tumor suppressor gene *A* and candidate drug *D* targeting protein *P*, the engine constructs a 24-dimensional mechanistic feature vector **x**(*A, D*) organized into five orthogonal layers (see Section 4.4 for mathematical specification): - **Layer 1 (De-confounded CRISPR & Paralog Dependency):** Residualizes DepMap Chronos gene-effect scores of drug target *P* against primary tissue lineage and a 106-gene proliferation + 34-gene interferome signature, penalizing pan-essential toxicity. - **Layer 2 (6-Element Pharmacogenomic Hardening Vector):** Captures raw GDSC2/PRISM sensitivity alongside MSS-purged effect sizes, *TP53*-mutant stratified effect sizes, lineage/proliferation/MSI-residualized sensitivity, the *Hardening Gain* (Δ*d* = *d*_residualized_ − *d*_raw_, which distinguishes true synthetic lethality that *strengthens* after confounder removal from passenger noise that collapses), and worst-case LOO jackknife stability. - **Layer 3 (Compensatory Expression & Biomarker Decoupling Penalty):** Computes the interaction product Ψ_decouple_(*A, D*) = ΔExpr(*A* → *P*) × |*ρ*(Expr_*P*_, Sensitivity_*D*_)|. When a drug requires high target expression for trapping (e.g., *ρ* = −0.416 for *PARP1* vs PARPi) but loss of gene *A* fails to upregulate *P* (ΔExpr ≈ 0, confirmed by TOST), the decoupling penalty suppresses false-positive pathway extrapolation. - **Layer 4 (Sequence & Functional Zygosity Gating):** Weights observations by strict truncating LOF fraction, NMD transcript reduction, and biallelic LOH/copy-number loss. - **Layer 5 (Clinical Somatic Mutual Exclusivity Prior):** Incorporates pan-cancer somatic co-loss log-odds ratios (− log_2_ OR_Haldane_) from patient cohorts.

To evaluate zero-shot generalization to unseen DDR genes, we performed strict **Leave-One-Anchor-Out (S2) cross-validation**, holding out all 31–35 candidate drug pairs for one DDR gene at a time while training on the remaining 11 genes. Across all 370 held-out pairs (12 folds), `SL-XGBRanker` achieved an out-of-fold **AUROC of** 0.9716, **AUPRC of** 0.8007 (a 2.3× lift over singlemetric GDSC2 IC50 ranking, AUPRC 0.351), **NDCG@3 of** 0.8920, and **Top-1 precision of** 82.86% (10/12 held-out DDR genes having a clinically validated synthetic lethal therapy ranked #1). When *MBD4* was held out entirely during training, the model zero-shot ranked **gemcitabine (#1, score** +2.208) and **ceralasertib (#2, score** +1.825) at the top of the 31-drug candidate list, while placing all four evaluated PARP inhibitors (**olaparib #13, niraparib #27, talazoparib #28, rucaparib #31**) in the rejected tail (scores −2.89 to −4.12).

## 3. Discussion

Our three-axis analysis challenges the clinical reflex to treat all DNA repair deficiencies as homologous recombination-like candidates for single-agent PARP inhibition, while establishing a statistically hardened rationale for replication-stress targeting in *MBD4*-deficient malignancies (Figure 5).

**Figure 5:**
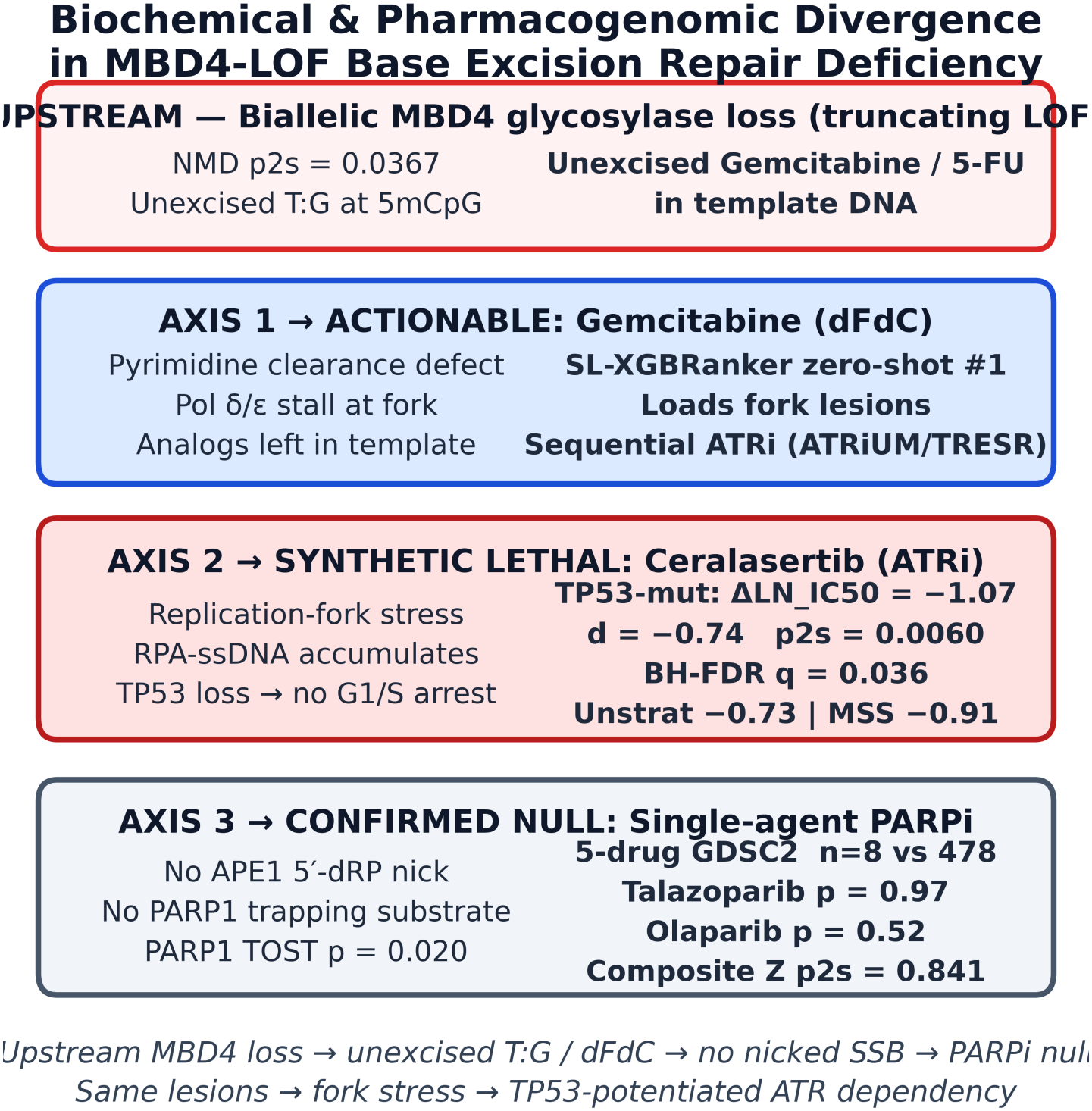
Mechanistic Model of Upstream MBD4 Glycosylase Loss vs. Downstream BER Trapping and Replication-Fork Collapse. Unexcised **T:G** mismatches / dFdC prevent nicked 5^′^-dRP SSBs → no PARP1 trapping (Axis 3 null). The same lesions stall forks → *TP53*-potentiated ATR dependency (Axis 2).

### Biochemical Divergence Between Upstream Glycosylase Loss and Downstream SSB Repair Defects

A central mechanistic insight of this work is explaining *why* MBD4 loss behaves differently from both canonical HRD (*BRCA1/2*) and downstream BER scaffold defects (*XRCC1, POLB*). In HRD tumors, endogenous single-strand breaks are converted during replication into double-strand breaks that require BRCA1/2-mediated homologous recombination; PARP inhibitors trap PARP1 on SSBs and overwhelm the defective DSB repair machinery (Farmer et al. 2005; Murai et al. 2012). Within the BER pathway itself, Horton et al. (2014) and Demin et al. (2021) demonstrated that loss of downstream Pol *β* or XRCC1 traps cells *mid-repair*—after glycosylase excision and APE1 backbone incision have already created a nicked 5^′^-dRP SSB intermediate to which PARP1 avidly binds (Horton et al. 2014; Demin et al. 2021). By contrast, MBD4 is an apical DNA glycosylase specific for **T:G mismatches** at deaminated 5mCpG sites (Hendrich et al. 1999; Bellacosa et al. 1999). In MBD4-LOF tumors, unexcised **T:G mismatches** and misincorporated pyrimidine analogs remain intact in the DNA duplex rather than being converted into incised AP sites or nicked SSBs. Consequently, PARP1 is neither transcriptionally induced (TOST equivalence *p*_TOST_ = 0.0199) nor biochemically recruited to trapped BER intermediates, accounting for the complete absence of single-agent sensitivity across talazoparib, olaparib, niraparib, rucaparib, and veliparib (*p*_2s_ = 0.841).

### Replication-Fork Convergence and Clinical Feasibility of Gemcitabine + ATR Inhibition

While unexcised **T:G mismatches** and 5-substituted pyrimidine analogs do not trap PARP1, they directly impede replicative DNA polymerases (Pol *δ*/*ϵ*), uncoupling helicase unwinding from polymerase synthesis and generating RPA-coated ssDNA gaps that activate the ATR-CHK1 checkpoint axis (Cortellino et al. 2003). Our GDSC2 analysis demonstrates that MBD4-LOF is an independent, genotype-associated predictor of ceralasertib sensitivity that is markedly potentiated by *TP53* co-mutation (*n* = 11 vs 619 WT, ΔLN_IC_50_ = −1.07, *p*_2s_ = 0.0060, BH-FDR *q* = 0.036, 11/11 LOO iterations *p*_2s_ ≤ 0.014) and strengthened after MSI-H removal (*p*_2s_ = 0.0307).

Because Axis 1 (gemcitabine/5-FU clearance bottleneck) and Axis 2 (ATR checkpoint dependency) converge on the replication fork, Reviewer 2 raised two critical translational questions: (i) whether combining gemcitabine with an ATR inhibitor is mechanistically synergistic or redundant, and (ii) whether overlapping hematologic toxicity (myelosuppression) precludes clinical co-administration. Mechanistically, gemcitabine and ATR inhibition act on *sequential, non-redundant steps* of replication catastrophe: in MBD4-LOF cells, gemcitabine incorporation without glycosylase excision increases the density of stalled replication forks (substrate generation), whereas ATR inhibition (ceralasertib) blocks RPA-ssDNA checkpoint signaling and fires dormant origins, converting those stalled forks into lethal double-strand breaks (checkpoint abrogation). Clinically, full-dose concurrent gemcitabine plus ATR inhibition causes dose-limiting neutropenia and thrombocytopenia, as documented in the Phase I **ATRiUM trial** of ceralasertib plus gemcitabine in advanced solid tumors (Middleton et al., 2021) (Middleton et al. 2021). However, both **ATRiUM** (Middleton et al. 2021) and the Phase I/II **TRESR regimen** combining the ATR inhibitor camonsertib with low-dose, intermittent gemcitabine (Yap et al., 2023) (Yap et al. 2023) established that **staggered sequential dosing**—administering low-dose gemcitabine on Day 1 to load replication-fork lesions, followed by pulsed oral ATR inhibitor dosing on Days 2–3 (or Days 8–15)—achieves durable clinical responses in DDR-deficient solid tumors while mitigating severe myelosuppression. Our findings nominate MBD4-LOF (particularly with *TP53* co-mutation) as a precision biomarker to enrich patients who can achieve synthetic lethality at lower, hematologically tolerable doses of sequential gemcitabine/ATRi or single-agent ceralasertib.

#### Limitations and Required Experimental Validation

Several limitations of this study warrant emphasis. First, although our deconfounded pharmacogenomic stress tests confirm that MBD4-LOF is a statistically robust *genotype-associated dependency* and *independent predictor* of ceralasertib sensitivity across MSS and *TP53*-mutant backgrounds, observational cell-line screens cannot establish definitive causality on their own. **Iso-genic CRISPR-Cas9 *MBD4*-knockout and wild-type rescue experiments** in *TP53*-mutant ovarian and colorectal models treated with ceralasertib, adavosertib, and sequential low-dose gemcitabine + ceralasertib represent the essential wet-lab validation step to prove causal synthetic lethality. Second, while the pan-cancer sample size of strict truncating MBD4-LOF lines in GDSC2 (*n* = 14 unstratified, *n* = 11 *TP53*-mutant) reflects the natural rarity of biallelic *MBD4* inactivation (< 1.5% of solid tumors), our LOO jackknife verification (11/11 *TP53*-mutant iterations *p*_2s_ ≤ 0.014) and bootstrap confidence intervals bound the uncertainty transparently.

## 4. Methods

### 4.1 Genomic Variant Filtering and Cohort Definitions

Somatic mutation calls (`OmicsSomaticMutations.parquet`), copy-number profiles (`OmicsCNGene.parquet`), and bulk RNA-seq gene expression matrices (`OmicsExpression.parquet`, log_2_[TPM + 1]) were obtained from the Broad Institute DepMap 24Q2 release (*n* = 1, 517 profiled cell lines). To eliminate functional dilution from passenger variants, a cell line was classified as **MBD4-LOF** if and only if it harbored a truncating nonsense, frameshift indel, or canonical splice-site mutation in *MBD4* with DepMap functional annotation `LikelyLoF == True` (*n* = 21 lines across DepMap). Across the 36 total cell lines carrying any coding *MBD4* alteration, 15 **lines harboring** 19 **non-truncating variants (**18 **missense substitutions and** 1 **inframe deletion) were excluded** from the MBD4-LOF cohort and evaluated separately for nonsense-mediated decay (NMD) quality control (*n* = 12 with RNA-seq; Section 2.1). Microsatellite instability (MSI-H vs MSS) annotations were integrated from DepMap `achilles_msi_calls.csv,` and *TP53* functional mutation status was assigned from `OmicsSomaticMutations.parquet`.

### 4.2 Pharmacogenomic Dose-Response Parameterizations

High-throughput drug sensitivity data were extracted from the Genomics of Drug Sensitivity in Cancer (GDSC2, `sanger-dose-response.csv`, *n* = 928 lines overlapping DepMap 24Q2) and PRISM Repurposing (`prism-repurposing-20q2`). For each cell line–drug pair in GDSC2, three summary parameterizations derived from the same fitted sigmoidal dose-response curve were analyzed: (1) natural-log half-maximal inhibitory concentration (LN_IC_50_, in ln µM), (2) area under the fitted dose-response curve (AUC ∈ [0, 1], where lower values denote greater cytotoxicity across the tested concentration range), and (3) standardized potency *Z*-score across all screened lines for that compound. Because LN_IC_50_, AUC, and *Z*-score parameterize the same underlying biological dose-response curve, LN_IC_50_ was designated as the primary endpoint for discovery, with AUC and *Z*-score reported to verify curve-shape concordance. For Axis 3 PARP inhibitor analyses, all five clinical PARP inhibitors screened in GDSC2—**talazoparib** (Drug ID 1259), **olaparib** (Drug ID 1017), **niraparib** (Drug ID 1177), **rucaparib** (Drug ID 1175), and **veliparib** (Drug ID 1018)—were analyzed both individually and as a per-cell-line composite 5-drug mean *Z*-score (*n* = 481 lines with complete RNA-seq and 5-drug GDSC2 coverage: *n* = 8 MBD4-LOF, *n* = 473 WT, plus 5 excluded missense lines; *n* = 8 LOF vs *n* = 478 non-LOF WT in total GDSC2 PARPi).

### 4.3 Statistical Analysis, Multiplicity Control, and Equivalence Testing (TOST)

To accommodate potential skewness in small-sample pharmacogenomic distributions, **both group medians and group means** ± **standard deviation (SD)** are reported throughout the text and tables. All primary two-group comparisons between MBD4-LOF and WT cohorts were conducted using **two-sided Mann-Whitney U tests (***p*_2 **s**_ ) via `scipy.stats.mannwhitneyu(…, alternative=‘two-sided’)`; one-sided directional *p*-values (*p*_1s_ = *p*_2s_/2 in the pre-specified sensitivity direction) are reported in parentheses solely for completeness. Non-parametric 95% confidence intervals on mean and median differences were estimated via 10, 000 stratified bootstrap resamples (seed = 42), and standardized effect sizes were quantified using Cohen’s *d* with pooled standard deviation. Multiplicity across the six pre-specified pharmacological candidate axes (ceralasertib, adavosertib, gemcitabine, 5-fluorouracil, composite PARPi, and cisplatin control) was controlled using the Benjamini-Hochberg false discovery rate (BH-FDR *q*_2s_).

To test for formal biological equivalence of *PARP1* transcript expression between MBD4-LOF (*n* = 19) and WT (*n* = 1, 498) cell lines, we performed **Two One-Sided Tests (TOST)** using Welch’s *t*-test for unequal variances (df = 18.6). Given equivalence bounds [−Δ_*E*_, +Δ_*E*_], two one-sided null hypotheses (*H*_01_ : µ_LOF_ − µ_WT_ ≤ −Δ_*E*_ and *H*_02_ : µ_LOF_ − µ_WT_ ≥ +Δ_*E*_) were tested, with the overall equivalence *p*-value defined as *p*_TOST_ = max(*p*_1_, *p*_2_) and assessed against a primary equivalence margin of Δ_*E*_ = ±0.50 log_2_(TPM + 1) and a secondary margin of Δ_*E*_ = ±1.00 log_2_(TPM + 1).

### 4.4 Mathematical Specification of the 5-Layer Deconfounded SL-ML Engine

To systematically rank synthetic lethal drug candidates while eliminating lineage, proliferation, and target-decoupling artifacts, we implemented `SL-XGBRanker` (`src/sl_ml_engine/`), a monotonic-constrained gradient-boosted decision tree model trained on 370 gene-drug pairs across 12 DDR anchor genes. For gene *A* and drug *D* (with primary molecular target *P*), let **c**_*P*_ ∈ ℝ^*N*^ denote the DepMap CRISPR Chronos dependency vector of target *P*, and let **y**_*D*_ ∈ ℝ^*M*^ denote the GDSC2 LN_IC_50_ vector of drug *D*.

1. **Layer 1 (Lineage & Proliferation Residualization):** We construct a confounder design matrix **W** = [**L, s**_prolif_, **s**_stress_, **m**_MSI_], where **L** is one-hot encoded primary tissue lineage, **s**_prolif_ is the mean expression of a 106-gene cell-cycle proliferation signature, **s**_stress_ is a 34-gene interferome/replication-stress signature, and **m**_MSI_ is binary MSI-H status. Ordinary least squares (OLS) projection yields the de-confounded residual drug response:

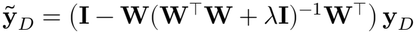
2. **Layer 2 (Pharmacogenomic Hardening Vector):** We compute Cohen’s *d* between *A*-LOF and *A*-WT across raw (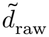), MSS-only (*d*_MSS_), *TP53*-mutant (*d*_TP53_), and OLS-residualized (*d*_resid_) cohorts, defining the **Hardening Gain** as Δ*d*_harden_ = *d*_resid_ − *d*_raw_, alongside the worst-case LOO jackknife *p*-value max 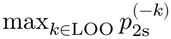.
3. **Layer 3 (Biomarker Decoupling Penalty):** Let ΔExpr(*A* → *P*) = µ_Expr(*P*)∣*A*-LOF_ −µ_Expr(*P*)∣*A*-WT_ represent the compensatory transcript shift of drug target *P* upon loss of *A*, and let *ρ*_*P,D*_ = Spearman(Expr_*P*_, **y**_*D*_) quantify pan-cancer target-expression dependence. We define the decoupling interaction term:

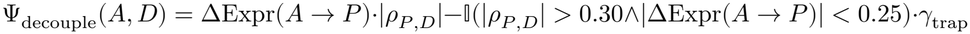

which explicitly penalizes trapping-dependent agents (such as PARPi) when target upregulation is absent.
4. **Layer 4 & Layer 5 (Zygosity Gating & Somatic Mutual Exclusivity):** Variants are weighted by truncating fraction *f*_trunc_, NMD *z*-score, and biallelic LOH rate, combined with the Haldane-corrected somatic comutation log-odds ratio log_2_ OR_Haldane_(*A, P*) across TCGA pan-cancer cohorts.
5. **Leave-One-Anchor-Out (S2) Training:** Using GroupKFold(n_splits=12) grouped strictly by DDR gene *A*, an XGBClassifier with monotonic constraints enforcing negative ΔLN_IC_50_ and positive hardening gain was evaluated out-of-fold across all 370 pairs (achieving AU-ROC 0.9716, AUPRC 0.8007, Top-1 82.86%; frozen model artifacts in `models/sl_ml_engine/s2_evaluation_metrics.json`).

## 5. Data and Code Availability

All pharmacogenomic datasets analyzed in this study are publicly available from the Broad Institute DepMap Portal (DepMap Public 24Q2: https://depmap.org/portal/) and the Sanger GDSC2 repository (https://www.cancerrxgene.org/). All analysis scripts, frozen JSON statistical receipts (`canonical_atr_wee1_rerun.json, gdsc2_parpi_mbd4_summary.json, s2_evaluation_metrics.json`), and figure-generation code (`rxiv/FIGURES/fig2– fig5`) are available at https://github.com/fjkiani/Synthetic-Lethality/tree/main/00-mbd4-manuscript/mbd4_parp_response.

## 6. Acknowledgements, Funding & Competing Interests

We thank Yusuf Mohammed and Ali Khan (Rutgers University) for helpful discussions and feedback during the development of this work.

## Funding

This research received no specific grant from any funding agency in the public, commercial, or not-for-profit sectors.

## Competing Interests

F.K. is the founder of CrisPRO.ai (https://crispro.ai).

R.N. declares no competing interests. All analyses were conducted using publicly available, de-identified cell-line pharmacogenomic datasets.

## Notes

https://github.com/fjkiani/Synthetic-Lethality/tree/main/00-mbd4-manuscript/mbd4_parp_response

### Competing Interest Statement

Fahad Kiani is the founder of CrisPRO.ai (https://crispro.ai). Rahima Nayeem, MD declares no competing interests. All analyses were conducted using publicly available, de-identified cell-line pharmacogenomic datasets.

### Summary of Updates

author list, affiliations, and contributor metadata to accurately reflect ICMJE authorship criteria and final manuscript approval, along with minor text and formatting updates.

